# *Sida cordifolia*, a medicinal plant, is efficacious in models of Huntington’s disease by reducing ER stress

**DOI:** 10.1101/2023.11.01.565080

**Authors:** Prasanna K Simha, Chandramouli Mukherjee, Karishma Bhatia, Vikas Kumar Gupta, Neerav M Sapariya, Padmanabhi Nagar, ZA Azim Nazeer, Ashwini Godbole, Bhavani Shankar Sahu, Sanjeev K Upadhyay

## Abstract

**Background and aim:** Huntington’s Disease is a severe neurodegenerative disorder caused by misfolded mutant huntingtin proteins with expanded stretches of polyglutamines aggregating and destroying cells in the nervous system. *Sida cordifolia* and *Acorus calamus* are medicinal plants used in traditional Ayurvedic medicine to treat neurological disorders. Here, we tested the effectiveness of extracts of both medicinal plants in decreasing aggregation of mutant huntingtin protein in models of Huntington’s Disease and explored the mode of action.

**Experimental procedure:** We used two models, the nematode *Caenorhabditis elegans* and a transgenic mouse neuroblastoma cell line, both expressing mutant huntingtin proteins with elongated polyglutamines. We assessed the effect of *Sida cordifolia* and *Acorus calamus* on mutant huntingtin protein aggregation in both models, and additionally used the cell line for mechanistic studies to identify cellular pathways underlying the effects of treatment.

**Results and conclusion:** Here, we show that an extract of *Sida cordifolia* inhibits aggregation of mutant huntingtin proteins. In the *C. elegans* model, the extract prolonged life span and improved motility of the nematode by reducing aggregation of the mutant huntingtin protein. *Acorus calamus* did not exhibit these effects. In the transgenic mouse neuroblastoma cell line, the extract decreased aggregation of the mutant huntingtin protein by suppressing key pathways in the ER stress response caused by the mutant protein. Our results highlight the potential therapeutic value of *Sida cordifolia* and its promise as a source for novel medications.

**Highlights:**

1. *Sida cordifolia* extract reduces aggregates in HD model of transgenic worms
2. Reduction in aggregates leads to improved motility and longevity
3. *Sida cordifolia* extract reduces ER stress in cells expressing mHTT protein
4. First report on the pharmacology of *Sida cordifolia* in neurodegeneration

## 1. INTRODUCTION

Huntington’s Disease (HD) is an autosomal dominant neurodegenerative disorder caused by the expansion of CAG trinucleotide repeats in the huntingtin (*Htt*) gene on chromosome 4. Longer CAG repeats cause earlier onset of disease and increased severity.^1^ The mutant genes encode huntingtin proteins with long tandem repeats of poly-glutamines (polyQ), typically more than 35 in a stretch. These misfolded mutant proteins aggregate and cause neurodegeneration.^1^ The disease is estimated to affect about 1 in 7500 individuals in the population^2^, being more prevalent in the Western population than in Asians.^1^ Cognitive, motor and psychological disturbances are attributed to neuronal degeneration in the striatum, globus pallidus, subthalamic nucleus, thalamus, substantia nigra and hypothalamus of the brain.^2^

Current treatments for HD are symptomatic and do not rectify the underlying causal cellular and molecular derangements. Symptomatic therapies include anti-dopaminergic agents and occasionally anti-depressants. Prolonged usage of these treatments in HD patients is associated with adverse effects^3^ including diarrhea, nausea, vomiting, depression, Parkinsonism and suicidal tendencies.^4,5^ In animal models, azo-dye congo-red^6^, disaccharide trehalose^7^, and polyQ-binding peptide-1^8^ have been reported to decrease the aggregation of mutant huntingtin protein, but none of these cross the blood-brain barrier, impeding their development as therapeutics. There remains an unmet need for new therapies to rectify the molecular derangements in HD, namely the aggregation of mutant huntingtin proteins and the toxic effects they cause in neurons.

Medicinal plants have been used in *Ayurveda*, an Indian traditional system of medicine, since the pharmacopoeia of approximately 70,000 formulations was developed over an extended period spanning 1500 BCE to the 18^th^ century CE. *Sida cordifolia* (Flannel Weed, Bala) (SC) is a perennially growing shrub of the mallow family Malvacea native to India. SC is used extensively in *Ayurveda* to treat diseases of the nervous system.

Ayurvedic formulations containing SC have been used to treat pain and neurological diseases associated with tremors or movement disorders^9^ and it has been found to be effective in Parkinson’s Disease^10^ and arthritis.^11^ Of interest, in one patient with early onset HD, treatment with a SC-containing Ayurvedic formulation, one of multiple formulations, caused a reduction in involuntary movements and improved balance and gait^12^. This case report inspired us to determine if SC could reduce aggregation of mutant huntingtin protein in models of HD. *Acorus calamus* (Sweet Flag, Muskrat root, Vacha) (AC), a plant from the family Acoraceae, is another neuroactive herb used in Ayurveda for epilepsy, speech disorders and enhancing intelligence.^13^ Both have been used safely in humans.

Here, we assessed the efficacy of both medicinal plants in HD. We used a *Caenorhabditis elegans* animal model and a mouse neuroblastoma cell line model. In *C. elegans* expressing a mutant huntingtin protein containing 40 tandem glutamines, a methanolic extract of SC, but not of AC, reduced aggregates of the polyQ-containing mutant huntingtin protein and thereby improved motility and prolonged life span of the nematode. In a transgenic mouse neuroblastoma cell line over-expressing a mutant huntingtin protein with 150 tandem glutamines, SC reduced aggregates of the polyQ-containing mutant protein by suppressing the Endoplasmic Reticulum Unfolded Protein Response (ER-UPR) triggered by aggregation. Since this extract likely has multiple active moieties, *Sida cordifolia* may be a source of novel therapeutics to inhibit aggregation of misfolded huntingtin proteins and the resulting neurodegenerative sequelae.

## 2. MATERIALS AND METHODS

### 2.1 Preparation of methanolic extract of *S. cordifolia* (*Bala*)

Roots of *Sida cordifolia* (*Bala*) were obtained from herb collectors in Tamil Nadu, India and authenticated by Prof. Nagar, Dept. of Botany, MSU, Baroda (Specimen no. SANJ20/1). The roots were washed thoroughly, air-dried, ground coarsely and used for cold maceration methanolic extraction. The roots were soaked in methanol (1:10 ratio, w/v) in a conical flask and incubated for 72h at 37°C in a shaker incubator. This mixture was filtered using Whatman filter paper. Rotatory vaporization was used to obtain the methanol-free extracts stored at -20°C for further use.

### 2.2 Preparation of methanolic extract of *A. calamus* (*Vacha*)

Rhizomes of AC were obtained from the botanical garden at Maharaja Sayajirao University (MSU), authenticated by Prof. Nagar, Dept. of Botany, MSU, Baroda (Specimen no. SANJ20/2). The rhizomes were thoroughly washed and dried, followed by powdering it coarsely. This plant rhizome powder was used for methanolic extraction (1:10 (w/v) root powder: methanol) using the soxhlet apparatus for 7-8h. The extract was concentrated and solvent was removed using a rotary evaporator and stored at -20°C for further use.

### 2.3 Culture and maintenance of *C. elegans*, a model of Huntington’s Disease

The *C. elegans* strains N2 (wild type) and AM141 (rmls133 (unc54p::Q40::YFP); expressing Q40::YFP in body wall muscles of the nematode were procured from the *Caenorhabditis* Genetics Centre (CGC), University of Minnesota, USA. The worms were grown at 20°C on Nematode Growth Medium (NGM) seeded with *Escherichia coli* OP50 using standard protocols. All worm experiments were conducted in accordance with the internationally accepted principles and approved by institutional committee (IBSC/TDU/05/12/22)

### 2.4 Culture of N2a neuroblastoma cells over-expressing a mutant huntingtin protein with expanded polyQ, and treatment with extracts of *S. cordifolia*

Transgenic mouse N2a cells expressing an ecdysone-inducible truncated N-terminal huntingtin gene with 150 CAG repeats attached to EGFP (HD 150Q) were a kind gift from Professor Nihar Jana’s group, National Brain Research Center, Manesar, India. They were used for evaluating the effects of the *S. cordifolia* extract on huntingtin aggregates. The cells between passage number 2 to 10 were used for all experiments. They were cultured in 10% (v/v) FBS in DMEM, with 0.4 mg/ml Zeocin and 0.4mg/ml G418 to its final concentration. For immunoblot experiments, 2 x10^5^ cells were plated in 6-well plates in replicates. Induction (with 1µM Ponasterone A) and treatment (1, 5, and 10µg/ml SC methanolic extract) were started simultaneously 24h after plating. Media was replaced with fresh media every 24h for two days, and after 48h of initial treatment, the cells were harvested and lysed for immunoblot or PCR (Figure 4A).

### 2.5 PolyQ aggregates – imaging

#### 2.5.1 In transgenic *C elegans*

Age-synchronized AM141 worms were grown up to Day 1 adults (72h at 20°C), on freshly prepared NGM-OP50 plates (60mm diameter) spotted with various concentrations (0.1µg/mL, 1µg/mL and 10µg/mL) of methanolic extracts of SC or AC. These worms were washed in M9 buffer and collected in sterile 1.5ml microfuge tubes. 20µL of 5mM sodium azide solution was added to the worm suspension to paralyze them. The worms were then mounted on glass slides containing a 2% agar pad and covered using a cover slip. The immobilized worms were observed under a fluorescent microscope (Olympus BX41) equipped with a camera (Olympus DP72). The images were captured under a blue filter (excitation wavelength 455nm to 495nm) using the Image-Pro Express software. The aggregates were defined as discrete structures with clear boundaries for counting puncta and quantified by measuring the average fluorescence intensity in the head region of the worm (up to the 2^nd^ pharyngeal bulb).

#### 2.5.2 In cells over-expressing mutant Huntingtin protein

2 x10^4^ cells were seeded in each well of 4 chamber slides, and after completion of treatment, cells were washed gently with PBS twice and then fixed using 4% PFA for 20 min. Thereafter, cells were mounted using DAPI containing mounting media (Fluorshield with DAPI SIGMA F6057-20ml) using 24mm*40mm cover glass (blue-star) and imaged on the next day in 40x oil Zeiss-Apotome microscope

### 2.6 Lifespan Assay

Age-synchronized AM141 worms were grown on freshly prepared NGM-OP50 plates (60mm diameter) spotted with various concentrations of preparation of *S. cordifolia* or *A. calamus*, up to late L4 stage. The worms were then transferred to fresh drug-containing NGM-OP50 plates every day until all worms in each group were dead. Worms were considered dead if they did not move after being lightly prodded using a platinum wire worm pick.

### 2.7 Thrashing Assay

Age-synchronized N2 or AM141 worms were grown on plates spotted with various concentrations of methanolic extracts of *S. cordifolia* or *A. calamus*, up to Day 1 adults (72h at 20°C). They were collected in M9 buffer and suspended in a 15µL droplet of the buffer on a glass slide. Thrashing of the worms was video recorded using a Levenhuk lite microscope camera. The thrashing was analyzed by counting the body bends manually. Same protocol was followed on day 3 and 5 of adulthood.

### 2.8 Immunoblotting

Following completion of treatment, cells were lysed with RIPA lysis buffer (50 mM Tris, pH 7.4, 150 mM NaCl, 1 mM EDTA, 0.1% SDS, 1% Triton X-100, 1% sodium deoxycholate with, 1mM phenylmethylsulphonyl fluoride, 2mM Na2VO5, 1 mM NaF, 20 mM Na4P2O7 as a phosphatase inhibitor cocktail) at 4°C. Lysates were collected, centrifuged at 14000xg for 15 min and the supernatant was collected. Total protein was estimated using the BCA (Bicinchoninic Acid) method. 20µg protein was loaded on SDS-PAGE and transferred to a nitrocellulose membrane. Membrane was blocked with 5% skimmed milk or 3% BSA (for phosphoprotein) for 2h at room temperature. Membranes were then probed overnight with primary antibodies of anti-GFP (rabbit,1:5000), anti-GAPDH (Mouse,1:8000), anti-pEIF2α (rabbit,1:2500), anti-EIF2α (rabbit,1:5000), anti- IRE1α (Rabit,1:5000), Anti pIRE1α (rabbit,1:2000), and washed with Tris-buffered saline, 0.1% Tween (TBST) and probed with HSP-tagged secondary antibody targeted against the host species of the primary (1:10000). Blots were developed using Luminol and chemiluminescence images were captured using the Uvitec Cambridge gel doc instrument.

### 2.9 Quantitative polymerase chain reaction (qPCR)

Total RNA was extracted using TAKARA RNAiso, following the manufacturer’s protocol. cDNA was prepared using BioRad iScript 5x RT supermix in a single-step reaction. qPCR was performed using iTaq SYBR green (BioRad) following the manufacturer’s protocol in CFX96 (BioRad). In another set of reactions, the synthesized cDNA was used to check the expression of the GFP transcript by semiquantitative PCR using GTPCR master mix (TAKARA BIO) for the GFP-specific primer, normalized with GAPDH. Primers are listed in table T1 in Supplementary Information.

## 3. RESULTS AND DISCUSSION

### 3.1 Methanolic extract of SC inhibits aggregation of polyQ-expanded mutant huntingtin protein in the *C. elegans* model of HD

The pathological hallmark of HD is the toxic aggregation of misfolded mutant huntingtin proteins with expanded tandem repeats of polyQ in the central nervous system. The nematode *C*. *elegans* is an excellent, widely established model organism for studying the properties of aggregated proteins and the resulting cellular effects.^14^ We investigated the therapeutic efficacy of the two neuroactive medicinal herbs, SC and AC, in the AM141 strain of *C. elegans* which expresses a mutant huntingtin protein with 40 tandem glutamines (Q40::YFP) in body wall muscles.^15^ These worms accumulate the polyQ protein aggregates as they age, which are seen as fluorescent punctae in the muscles (Fig 1A). We cultured these worms on fresh NGM/OP50 plates spotted with methanolic extracts of SC or AC. Worms treated with the SC extract, from embryo stage to day 1 adult stage (72h at 20°C), showed about 20-30% reduction in the number of YFP puncta (a measure of aggregated mutant huntingtin protein) as compared to vehicle/solvent (DMSO)-treated worms (Fig. 1A, 1B and S1). Maximum reduction was seen at 1μg/ml of SC extract, and the effect diminished upon treatment with higher or lower concentrations. The average fluorescence levels of the puncta were also lower upon SC treatment (Fig. 1C). This effect was specific to the SC extract, as the methanolic extract of AC did not reduce mutant huntingtin protein aggregation.

**Fig. 1:**
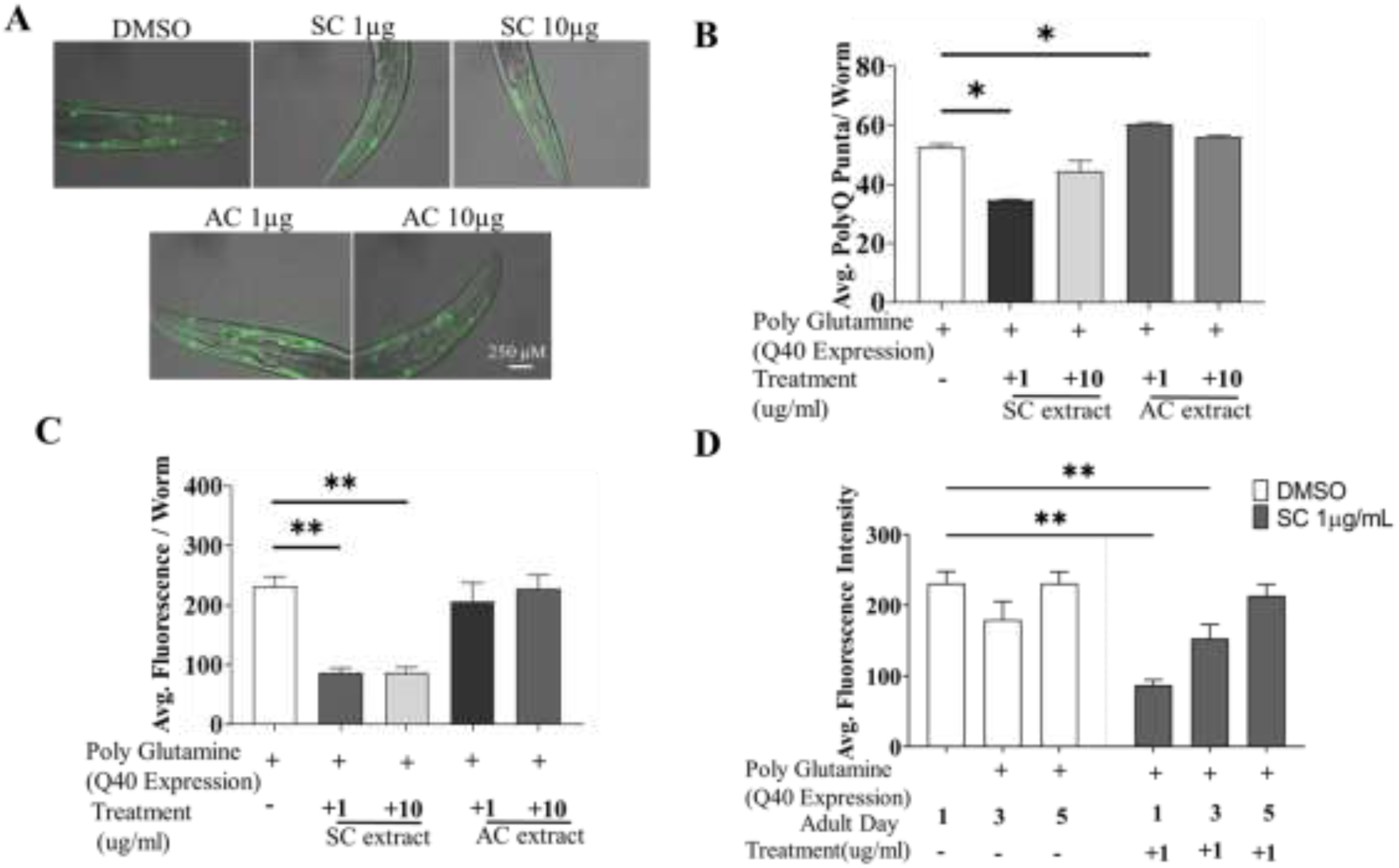
SC reduces polyQ aggregates in transgenic HD model of C *eleguns* (AM141). Poly Q aggregation in AML41 worms treated with SC or AC extracts (A) Representative fluorescent microscopy images (day 1 adult worms) (B) number of poly-Q punctac per worm (day 1) (C) average fluorescence intensity of poly-Q per worm (D) fluorescence intensity of poly-Q in Day 1, 3 and 5 adult worms. Data arc mean + SD, *p<0.05, **p<0.01

We next performed imaging-based quantification of polyQ aggregates in worms at days 1, 3 and 5 of adulthood. Worms treated with 1µg/ml SC significantly reduced fluorescence intensity of polyQ-YFP on day 1, and this effect gradually dissipated by day 5 (Fig.1D).

Taken together, these results show that the SC extract (1µg/ml) reduces aggregation of the mutant polyQ huntingtin protein.

### 3.2 SC treatment improves motility in the *C. elegans* model of HD

*C. elegans* swim by producing alternating C-shape and straight conformations called “thrashing”.^16^ Aggregation of polyQ mutant huntingtin proteins in the body wall muscle leads to defects in thrashing. We performed the thrashing assay to investigate whether SC treatment improved motility of HD worms through its reduction of aggregated mutant huntingtin proteins. Worms grown on fresh NGM/OP50 plates containing different concentrations of SC or AC extracts were video graphed and their motility checked with the thrashing assay on days 1, 3 & 5 of adulthood. Treatment with SC (1μg/mL), but not AC, restored motility of mutant worms to that of wild-type worms on day 1 (Fig. 2A and B), the effect decreasing by day 5 (Fig. 2C and 2D, S2).

**Fig. 2:**
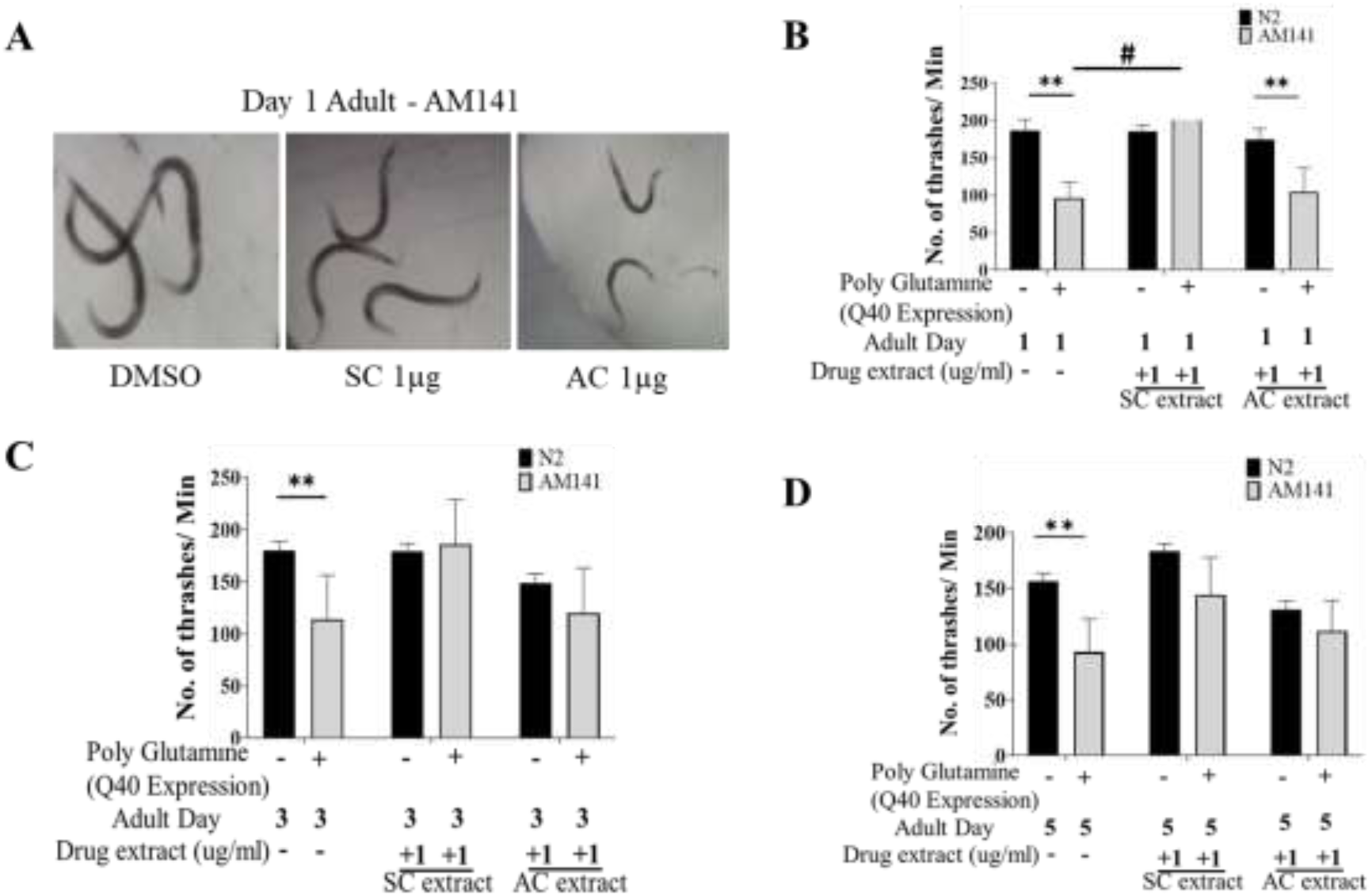
SC improves motility in HD model of G *elegans.* Thrashing assay of N2 and AM141 worms treated with SC or AC extracts. (A) Representative images of thrashing of Day I adult AVI 141 worms. Quantification of thrashing of (B) Day 1. (C) Day 3 and (D) Day 5 adult AM141 worms. **p<0.01, #p<0.0001. Data represented as mean ± SD.

### 3.3 SC treatment extends life span of HD model of *C. elegans*

Earlier studies showed that lifespan extension in *C. elegans* is neuroprotective.^15^ We investigated whether SC would extend the lifespan of *C. elegans*. Worms were treated with differentconcentrations of SC or AC extract on NGM OP50 plates from the embryo stage. SC extended the life span of AM141 worms expressing the 40Q mutant huntingtin protein, from 11.55 days in untreated worms to 17.07 days in SC-treated (1µg/mL) worms (3A and 3D). In contrast, treatment with AC reduced life span of these worms (Fig. 3B, C and D).

**Fig 3:**
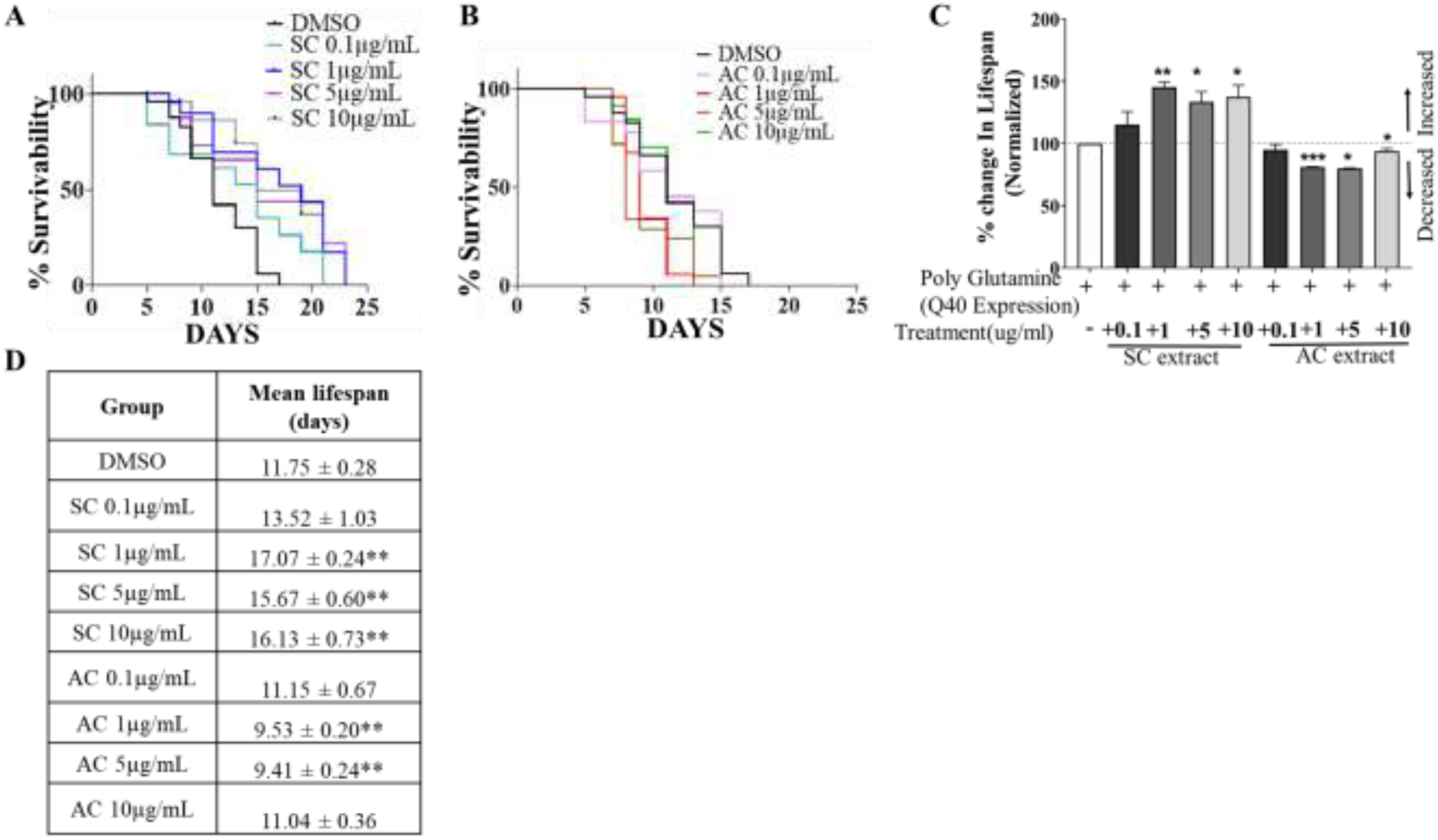
SC improves lifespan of HD model of *C. elegana.* Survival curves of AM 141 worms (A) treated with SC (B) treated with AC (C) Quantification of percentage change in lifespan of AM141 worms treated with SC or AC extracts (D) Mean change in survival. n=25 per group. Data represented as mean ± SD. **p<0.0l

### 3.4 SC reduces aggregation of mutant polyQ-containing huntingtin proteins in a mammalian system by reducing ER stress

We tested if SC could suppress aggregation of mutant huntingtin proteins in a mammalian system. For these studies we used the mouse transgenic N2a neuroblastoma cell line over- expressing an ecdysone-inducible mutant Huntingtin protein with an expanded polyQ repeat (150Q) attached to EGFP (HD150Q-EGFP).^17^ Upon treatment of these cells with 1µM Ponasterone A, analog of Ecdysone (Fig. 4A), the HD150Q-EGFP mutant protein aggregated, visualized as increased GFP puncta by fluorescence microscopy. SC treatment significantly reduced HD150Q-EGFP aggregates (Fig. 4B and 4C) by decreasing mutant protein levels (Fig. 4E and 4F) without altering mRNA levels (transcription) (Fig 4D).

We next determined if SC-mediated reduction of mutant huntingtin protein aggregation was due to modulation of the Endoplasmic Reticulum Unfolded Protein Response (ER-UPR). In mammals, three main pathways initiated by three different type-1 transmembrane proteins: inositol – requiring protein – 1α (IRE -1α), protein kinase RNA-like ER kinase (PERK) and activating transcription factor-6 (ATF6), govern ER-UPR. XBP-1, a transcription factor critical in this process, is induced by ATF6, spliced by IRE-1, and modulating XBP-1 is beneficial in animal models of HD.^18^

Ecdysone-induced aggregation of HD150Q-EGFP protein triggered ER-UPR as evidenced by increased phosphorylation of IRE-1α (Figs 5A, B), splicing of XBP-1 (Fig 5C) and activation of phosphor-PERK-dependent phosphorylation of eIF2A (Fig 5D, E). SC treatment suppressed all these changes (Figs 5A, B, C, D and E).

**Fig 4:**
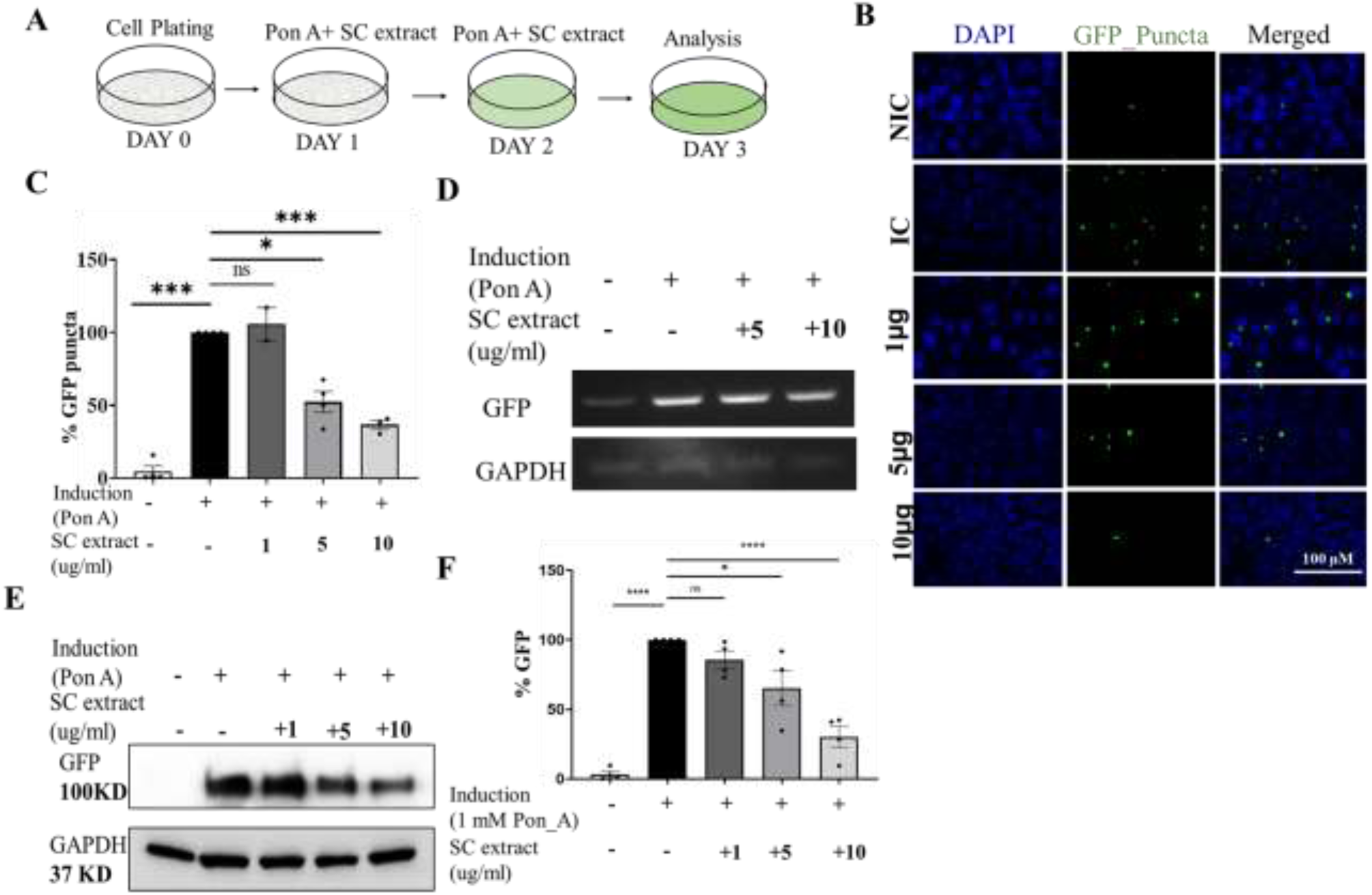
SC reduces protein aggregate in the poly-Q-EGFP expressing cells. (A) Treatment protocol (B) Representative images of SC treated cells (C) Quantification of GFP- puncta per cell. (D) Semi-quantitative PCR of GFP mRNA expression (E) Immunoblot of GFP protein (F) Quantification of immunoblots of GFP. All data represented as mean ± SEM, *p<0.05,**p<0.01, ***p<0.001 and ns is statistically non-significant.

**Fig 5:**
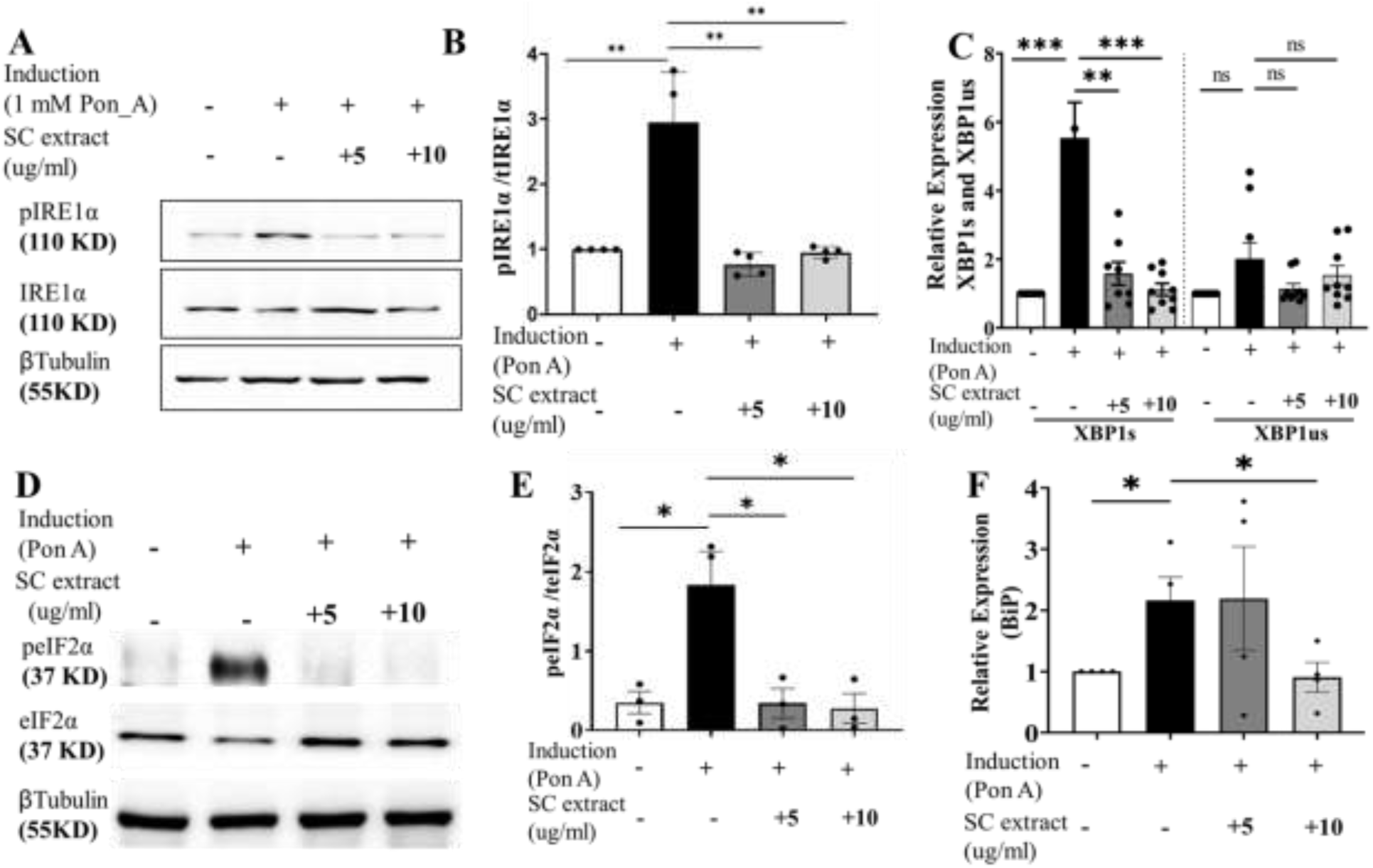
SC modulates ER stress pathway. (A) Immunoblots of plrelα and total Irei α (B) and its quantification (C) Relative expression of *xbpls* and *xbplus* (D) Representative Immunoblot of pEif2α and total Eif2α (E) its quantification (F) Relative expression of *bip.* All data represented as mean±SEM, *p<0 05,**p<0.01, ****p<0.001 and ns is statistically non-significant.

SC also suppressed other ER stress responses induced by HD150Q-EGFP protein aggregation including reduction of transcript levels of ATF4 and CHOP (Fig S3). CHOP, a protein linked to apoptosis,^19^ is a key link between the ER stress pathway and apoptosis. SC-mediated reduction in CHOP raises the possibility that SC may inhibit apoptosis in stressed cells as a feature of neuroprotection. Protein levels of BiP, an important chaperone in the ER stress response, were also reduced by SC treatment (Fig. 5F).

## Conclusion

In summary, SC treatment reduces the aggregation of mutant polyQ-containing huntingtin proteins both in a *C elegans* animal model and in a mouse neuroblastoma cell line (Fig 6). These effects are, at least in part, due to the suppression of various pathways in the ER stress response (Fig. 6). Previously described experimental medications that reduced aggregation of mutant huntingtin protein in animal models (azo-dye congo-red^6^, disaccharide trehalose^7^, and polyQ-binding peptide-1^8^) had to be administered directly into the central nervous system because they were not able to penetrate the blood-brain-barrier. In contrast, the SC extract was effective in reducing aggregation of mutant huntingin protein in *C. elegans* when applied into the bathing medium, suggesting that it readily crosses the blood-brain-barrier. Further, oral administration of a SC-containing Ayurvedic formulation to a patient with juvenile-onset HD improved motor function, balance and gait without any adverse effects^12^, again suggesting that SC-containing formulations can cross the blood-brain barrier.

**Fig 6:**
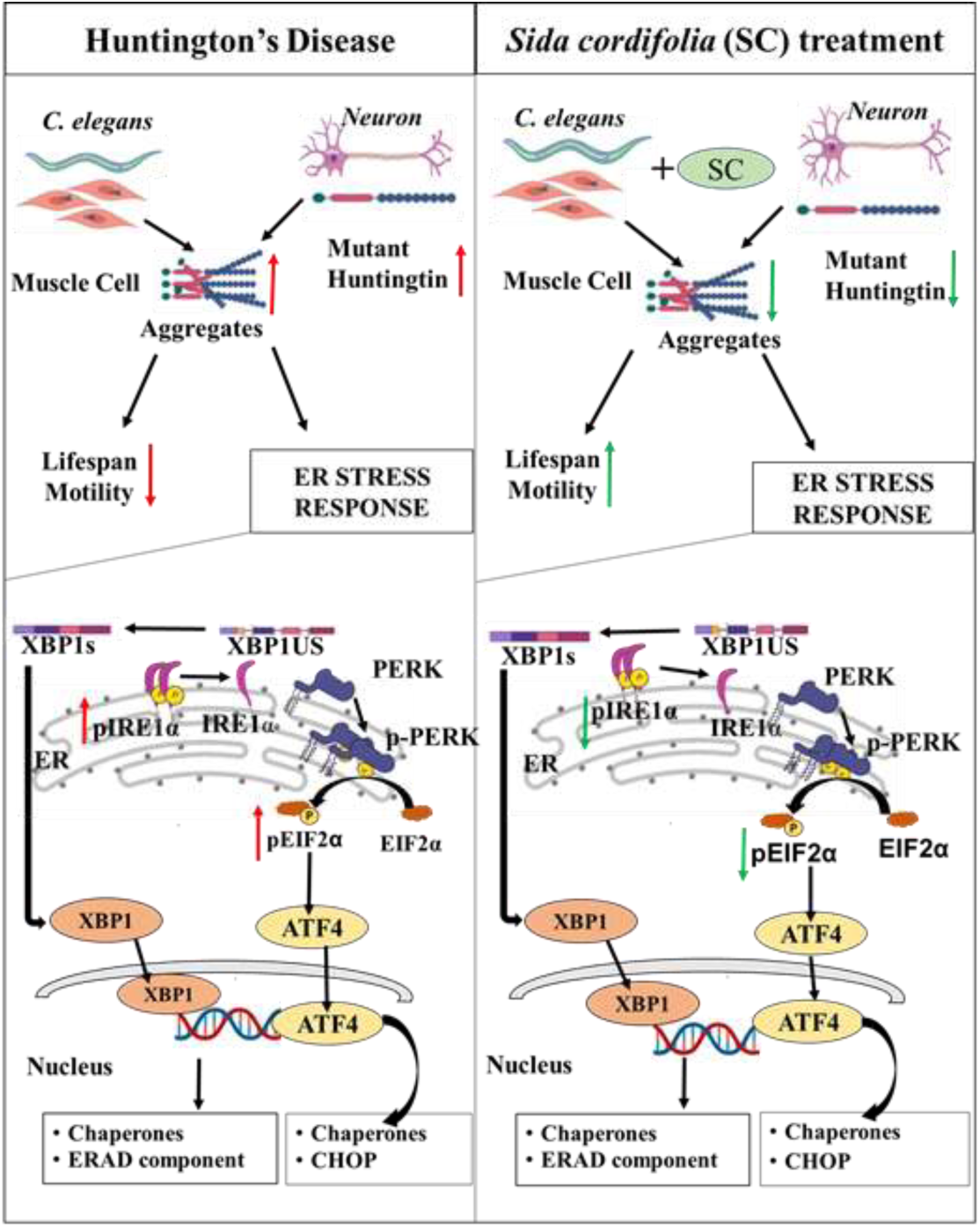
*Sida cordifolia* is efficacious in HD models by reducing ER stress. reduces the aggregates in transgenic HD worm model expressing mutant protein (Q40::YFP) in their body wall muscle. This improves motility and enhances the lifespan of these worms. In neuroblastoma cells (N2a) expressing polyQ- EGFP SC treatment reduces the amount of mutant protein and thereby reducing the number of aggregates. This is accompanied by reduction in levels of key ER stress markers - Spliced XBPls, pEIF2α and pIRElα.

The methanol extract of SC likely contains several active ingredients that modulate multiple molecular targets. Future studies using fractionation of the extract coupled with cell and animal-based studies may help to identify active moieties within the extract and the molecular targets they modulate. Such an approach may also lead to the discovery of novel therapeutics that correct the molecular derangements in HD. Our study highlights the importance of applying modern molecular and cell biology approaches to evaluate medicinal plants and identify new drug entities from scientifically validated botanical products.^22^

## Abbreviations

SC: Sida cordifolia
AC: Acorus Calamus
HD: Huntington’s Disease
mHTT: mutant (truncated) Huntingtin protein

## Acknowledgement

We acknowledge the help extended by Mr. Anjaneyulu J, in collection and preparation of the SC extract. We thank Prof. George Chandy for providing valuable inputs to the project.

## Fundings

The authors thank UGC for a start-up grant (F.4-5(185-FRP)/2015) to SKU and Pratiksha Trust for a research grant to AG. We also thank CSIR for fellowship to KB and ICMR for fellowship to VKG. Ramalingaswami fellowship grant (BT/RLF/Re-entry/38/2016) and core funds from NBRC funded the BSS lab towards this study. We thank NBRC core funds for providing salary support to BSS and CM.

## Ethical Statement

All worm experiments were conducted in accordance with the internationally accepted principles and approved by institutional committee of TDU (IBSC/TDU/05/12/22 dated Dec 5th, 2022).

## Data availability

We have provided all relevant data in the forms of figures and supplementary materials. Any additional data will be provided on request.

---

**Supplementary Figure S1:**
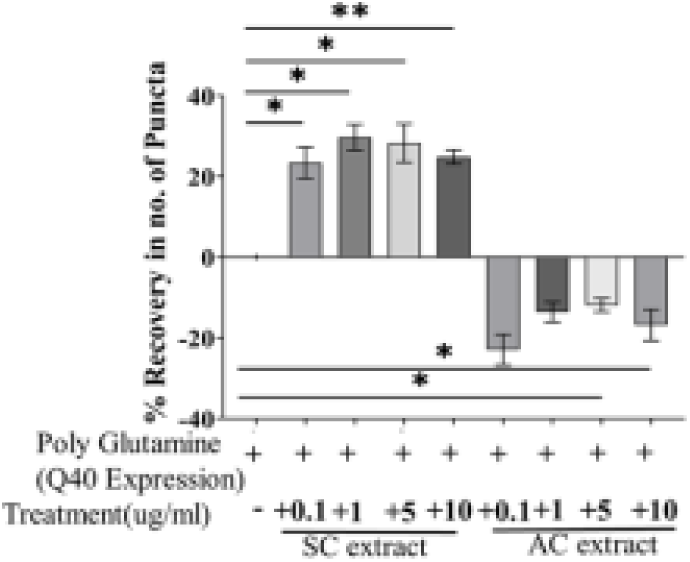
SC reduces polyQ aggregates in HD model of *C. elegant.* Percentage recover data of no. of puncta in Day I adult worms treated with SC or AC extracts. •p<0.05**p<0.01. Data arc mean ±SD.

**Supplementary Figure S2:**
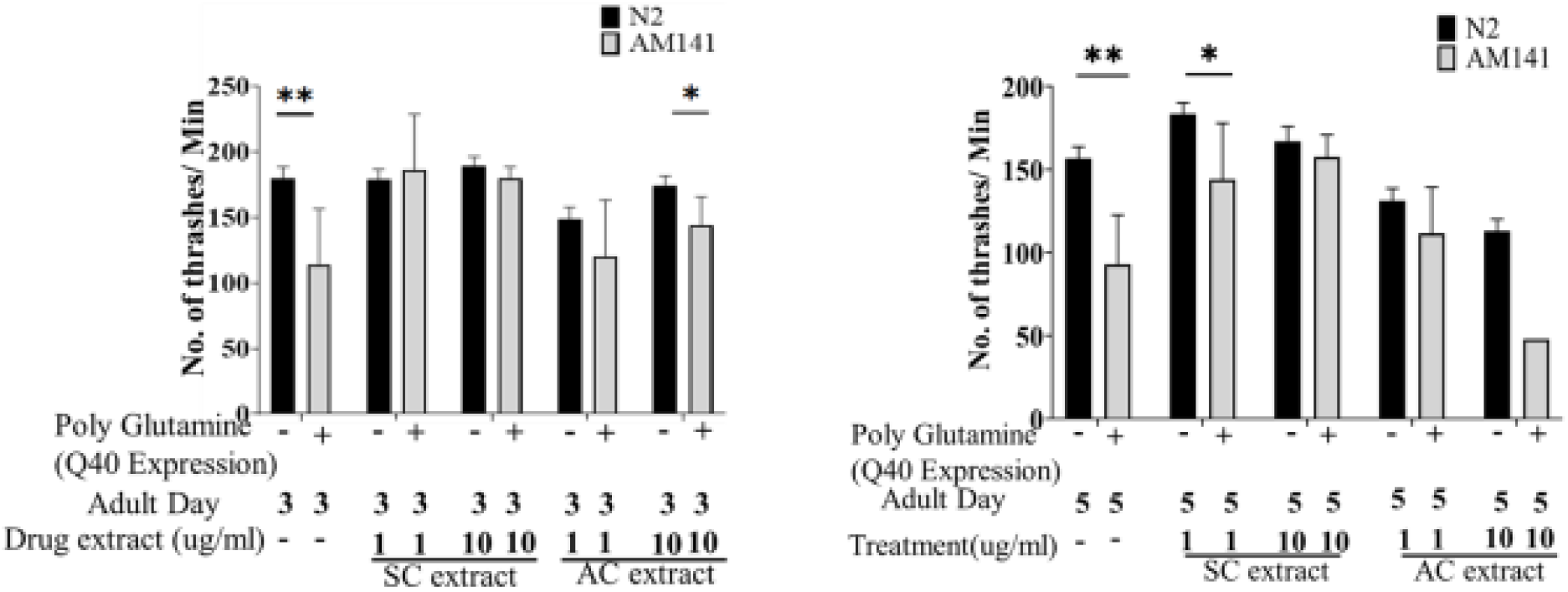
SC improves motility in HD worms. Quantification of thrashing assay of day 3 (A) and day 5 (B) adult AMI 41 and N2 worms treated with SC or AC extract. *P<0.05, **p<0,01. data arc mean ±SD

**Supplementary Figure S3:**
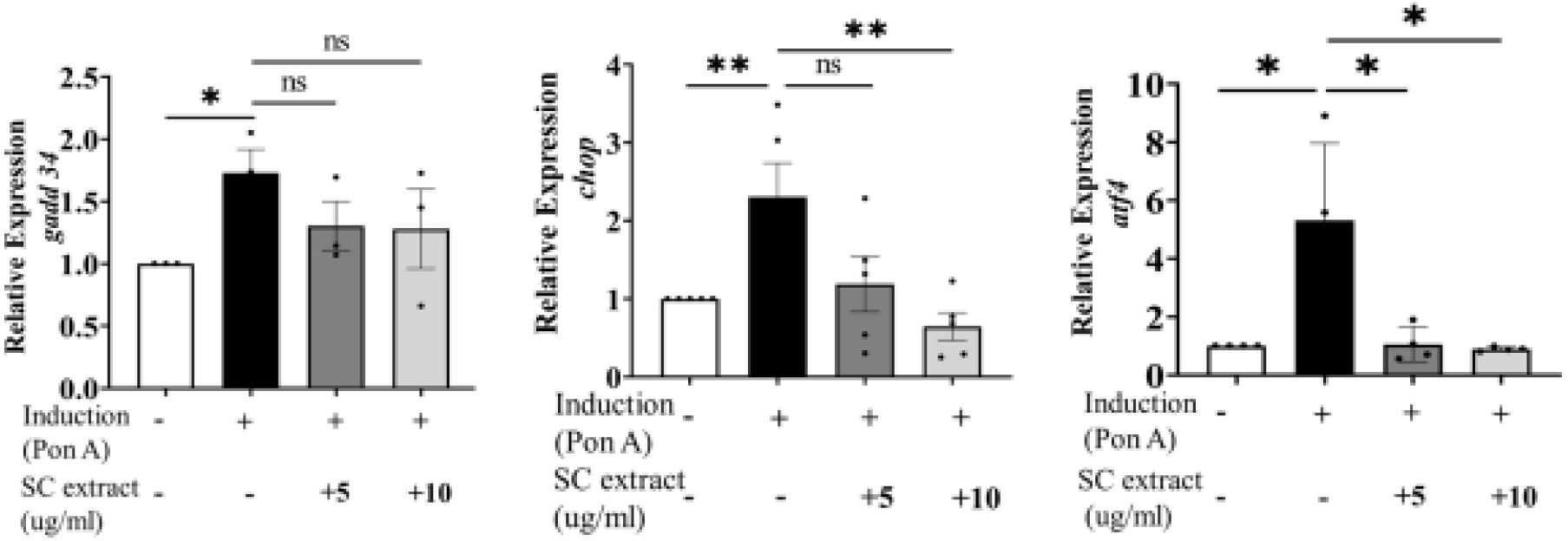
Relative expression GADD34 (A), CHOP(B), and ATF4 (C) in polyQ-EGFP expressing N2a cells. *P<0.05, **p<0.0l and ns non significant. Data are mean ±SD

**S4.**
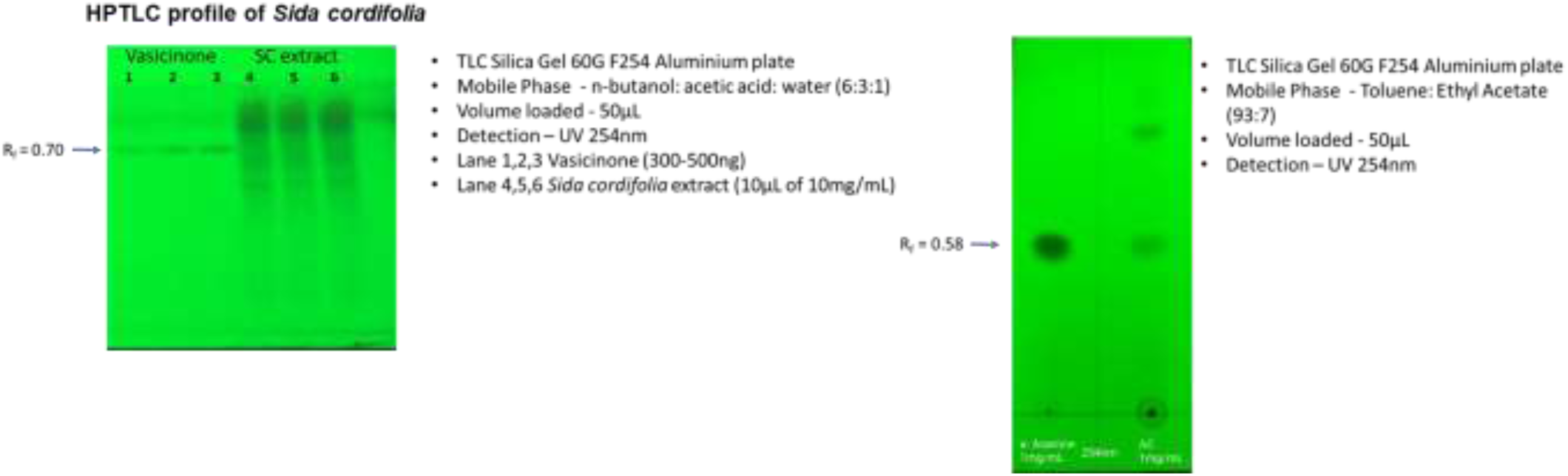

**Table T1:**
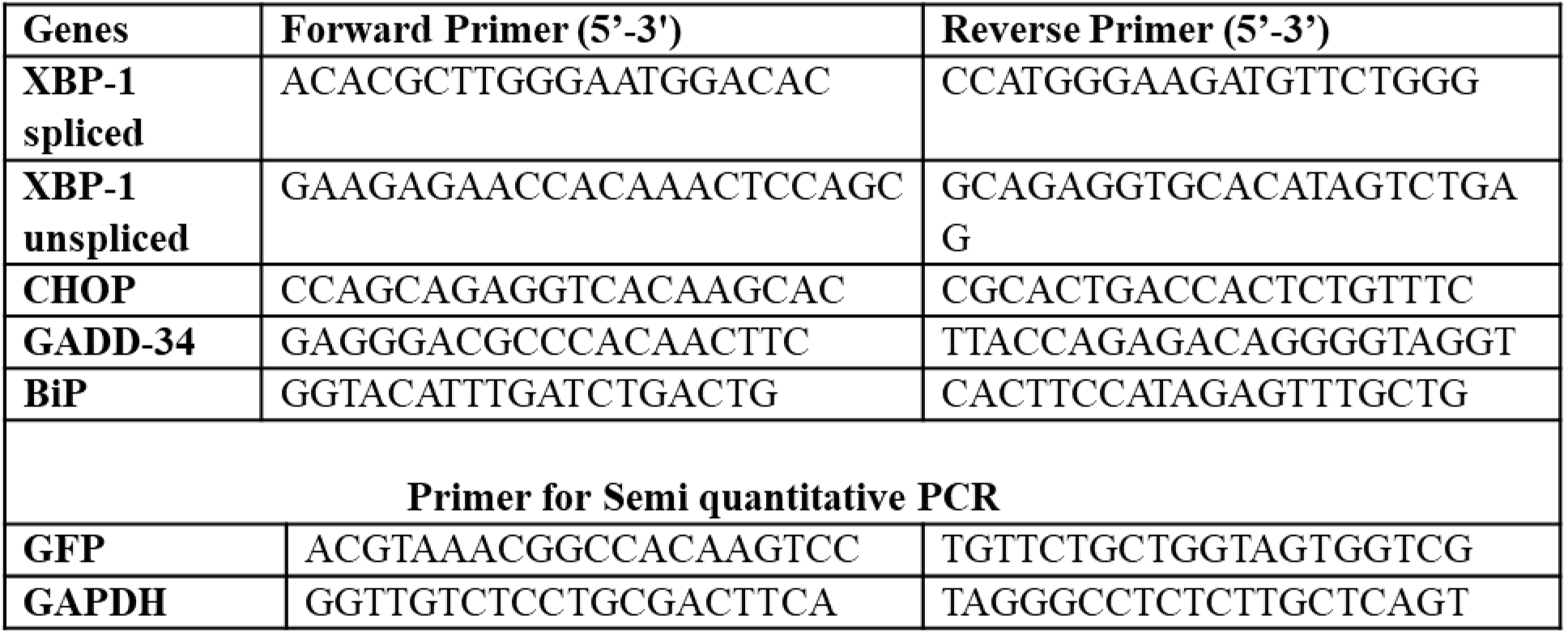
Primers used in the study.

## Notes

**Declaration of competing interest:** The authors declare that they have no known competing financial interests or personal relationships that could have appeared to influence the work reported in this paper.

### Competing Interest Statement

The authors have declared no competing interest.

### Summary of Updates

Figure number 1 of the previous version has been inadvertently displayed twice due to a formatting issue which was noticed after the screening board accepted the pre-print. We would like to correct this and if possible remove the previous version

